# Shortening of 3’ UTRs in most cell types composing tumor tissues implicates alternative polyadenylation in protein metabolism

**DOI:** 10.1101/2021.06.30.450496

**Authors:** Dominik Burri, Mihaela Zavolan

## Abstract

During pre-mRNA maturation 3’ end processing can occur at different polyadenylation sites in the 3’ untranslated region (3’ UTR) to give rise to transcript isoforms that differ in the length of their 3’UTRs. Longer 3’ UTRs contain additional cis-regulatory elements that impact the fate of the transcript and/or of the resulting protein.

Extensive alternative polyadenylation (APA) has been observed in cancers, but the mechanisms and roles remain elusive. In particular, it is unclear whether the APA occurs in the malignant cells or in other cell types that infiltrate the tumor. To resolve this, we developed a computational method, called SCUREL, that quantifies changes in 3’UTR length between groups of cells, including cells of the same type originating from tumor and control tissue. We used this method to study APA in human lung adenocarcinoma (LUAD).

SCUREL relies solely on annotated 3’UTRs and on control systems, such as T cell activation and spermatogenesis gives qualitatively similar results at much greater sensitivity compared to the previously published scAPA method.

In the LUAD samples, we find a general trend towards 3’UTR shortening not only in cancer cells compared to the cell type of origin, but also when comparing other cell types from the tumor vs. the control tissue environment. However, we also find high variability in the individual targets between patients. The findings help to understand the extent and impact of APA in LUAD, which may support improvements in diagnosis and treatment.

## Introduction

The processing of most human pre-mRNAs involves 3’ end cleavage and addition of a polyadenosine (poly(A)) tail. Typically, there are multiple cleavage and polyadenylation sites within a gene, and alternative polyadenylation (APA) has emerged as a major source of transcriptome diversity (Reyes & Huber, 2018). A prevalent type of APA isoforms are those that differ only in the length of their 3’ untranslated regions (3’ UTRs). 3’ UTRs become shorter upon T cell activation (A. R. Gruber et al., 2014; Sandberg et al., 2008), in cancer cells (Mayr & Bartel, 2009; Xia et al., 2014) and upon induction of reprogramming in somatic cells (Ji & Tian, 2009). Although the responsible regulators are still to be determined, core 3’ end processing factors under the transcriptional control of cell cycle-related transcription factors have been implicated, at least in the context of cell proliferation (Elkon et al., 2012). Various RNA-binding proteins (RBPs) are also involved in specific cellular systems (A. J. Gruber, Schmidt, et al., 2018; Lee et al., 2021; Martin et al., 2012; Masuda et al., 2020; So et al., 2019).

While APA-dependent 3’ UTR shortening has been observed in many cancers (Schmidt et al., 2018; Xia et al., 2014), it is presently unclear whether it is a manifestation of the change in cell composition of the tissue or of functional changes in all cell types within the tumor environment. As single cell RNA sequencing (scRNA-seq) technologies specifically capture mRNA 3’ ends, and datasets of tumor and matched control tissue samples have started to become available, this question can now be addressed, provided a few challenges are overcome. First, the number of transcripts that can be reliably quantified is still low (Breda et al., 2021), because the total number of reads obtained from individual cells is in the 10^3^-10^4^ range. Thus, quantifying gene expression at the isoform level is still very challenging. This issue can be partially circumvented by pooling the reads from cells of the same type. Second, while 3’ biased, scRNA-seq reads do not always reach the PAS and may also result from internal priming. Thus, identifying which reads correspond to the same 3’ end is also not trivial. This problem can be mitigated by associating scRNA-seq reads with already-annotated transcript 3’ ends. However, the current annotation is still far from complete (A. J. Gruber, Gypas, et al., 2018), leading to PAS usage quantification that is imprecise and incomplete. For this reason we developed a PAS-agnostic approach for quantifying changes in 3’ UTR length between samples, based on the entire 3’ end read distribution along the 3’ UTR. Applying the method to single cell sequencing data from human lung adenocarcinoma (LUAD), we found that 3’ UTR shortening is not specific to a cell type but rather occurs in most cell types that compose the tumor. Furthermore, our analysis revealed that the targeted transcripts encode proteins that are involved in various steps of protein metabolism, including synthesis at the endoplasmic reticulum (ER), transport between ER and the Golgi network and finally secretion of proteins. Our data thus implicates APA in the remodeling of protein metabolism in tumors.

## Results

### A myeloid to lymphoid switch in lung tumors

While analyses of bulk RNA-seq data revealed the shortening of 3’ UTRs in virtually all studied cancers with respect to matched control tissue, the shortening is especially pronounced in lung tumors (A. J. Gruber, Schmidt, et al., 2018). Thus, to better understand the mechanism and function of APA in cancers, we identified two studies in which single cell sequencing of lung adenocarcinoma (LUAD) and matched control tissue from multiple patients was carried out on the same platform, 10X Genomics (Lambrechts et al., 2018; Laughney et al., 2020). These data enable us to not only identify 3’ UTR changes in specific cell types, but also to assess their generality between studies and patients. We followed the procedure described in (Lambrechts et al., 2018) to annotate the type of individual cells. Briefly, we integrated the data with the *harmony* package (see Methods, Suppl. Fig. 1), clustered the normalized gene expression vectors of all cells (Fig. 1A) with the *Seurat* package (Butler et al., 2018), and annotated the type of 38’156 cells from 12 samples of the (Lambrechts et al., 2018) study (samples 3a-d, 4a-d, 6a-d, representing 3 tumor samples and a matched control for each of three patients) and 18’543 cells of the (Laughney et al., 2020) study (3 pairs of tumor-matched control samples) based on known markers. We used the markers proposed in the (Lambrechts et al., 2018) study, but also added a few markers for mast cell (*TPSAB1, TPSB2* and *CPA3*; (Dwyer et al., 2016) Table 1) (Fig. 1B). As described in the initial study (Lambrechts et al., 2018), the most abundant cell types in the tumor samples were T cells, myeloid and B cells, while the matched control samples were dominated by myeloid and alveolar cells (Fig.1C). We further identified a small cluster of mast cells, annotated as B cells in the initial study that did not consider mast cell markers. We observed a similar myeloid to T cell switch between control and cancer samples from the (Laughney et al., 2020) study (Fig. 1D). In addition, the matched control samples from this latter study had a more homogenous cell type composition compared to those from the (Lambrechts et al., 2018) study, consisting almost exclusively of lymphocytes and myeloid cells (Fig. 1D).

**Figure 1.**
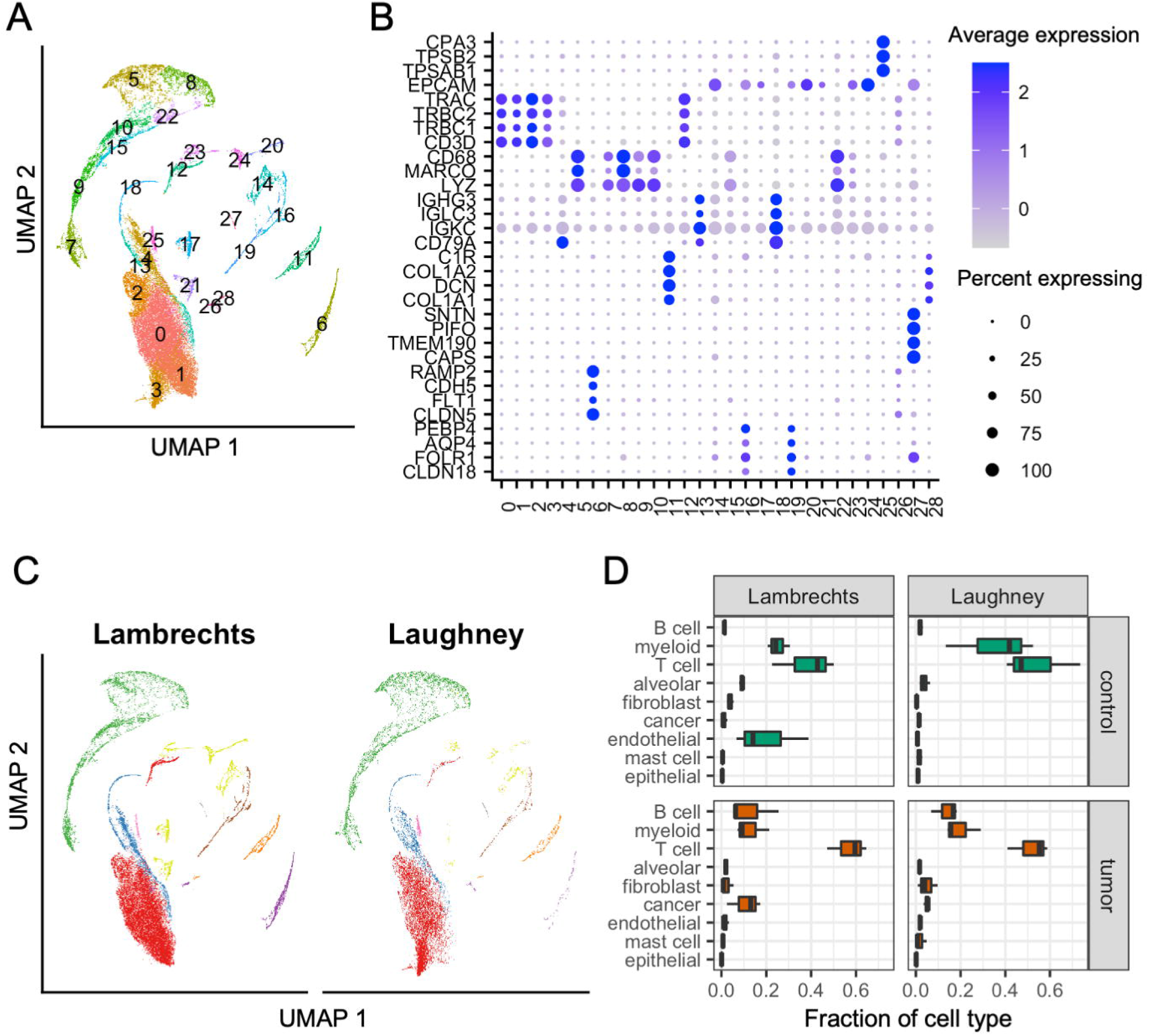
Cell type composition of lung adenocarcinoma and matched control samples. **A.** 2-dimensional projection (Uniform Manifold Approximation and Projection, UMAP) of gene expression vectors. The projections were obtained with the *RunUMAP* function from *Seurat* v3.2.3 (Butler et al., 2018), based on the first 10 principal components. The two datasets were integrated with *harmony*. Cell clustering was done on the shared nearest neighbour (SNN) graph (see Methods). **B.** Dot plot of marker gene expression across the clusters shown in panel A. Shown is the average expression and percent of expressing cells per cluster for the markers used in (Lambrechts et al., 2018) (see also Table 1). The dot plot was created with *Seurat*. **C.** 2-dimensional projection (created with *Seurat*) of gene expression vectors as in A, but highlighting only cells from one study in each panel. **D.** Box plot of relative proportion of each cell type in control (green) and tumor (red) samples from individual patients from the Lambrechts and Laughney datasets.

**Table 1:** Marker genes for cell type annotation. Based on (Lambrechts et al., 2018).

| Cell type | marker genes | Cell type | marker genes |
| --- | --- | --- | --- |
| alveolar | *CLDN18* | fibroblast | *C1R* |
| alveolar | *FOLR1* | B cell | *CD79A* |
| alveolar | *AQP4* | B cell | *IGKC* |
| alveolar | *PEBP4* | B cell | *IGLC3* |
| endothelial | *CLDN5* | B cell | *IGHG3* |
| endothelial | *FLT1* | myeloid | *LYZ* |
| endothelial | *CDH5* | myeloid | *MARCO* |
| endothelial | *RAMP2* | myeloid | *CD68* |
| epithelial | *CAPS* | myeloid | *FSGR3A* |
| epithelial | *TMEM190* | T cell | *CD3D* |
| epithelial | *PIFO* | T cell | *TRBC1* |
| epithelial | *SNTN* | T cell | *TRBC2* |
| fibroblast | *COL1A1* | T cell | *TRAC* |
| fibroblast | *DCN* | cancer | *EPCAM* |
| fibroblast | *COL1A2* |  |  |

Given that T cells are the most numerous cell type in tumor samples and that T cell activation leads to 3’ UTR shortening (A. R. Gruber et al., 2014; Sandberg et al., 2008) we wondered whether the pattern of 3’ UTR usage that was previously inferred from ‘bulk’ samples can be attributed to the infiltration of the tumor with activated T cells. To investigate this possibility, we first determined the distribution of RNA molecules (unique molecular identifiers, UMI) per cell in various cell types in the two studies (Suppl. Fig. 2A) and the total number of UMIs obtained from each cell type in each data set (Suppl. Fig. 2B). While T cells were the most numerous cell type in tumors, their relatively small RNA content per cell led to a smaller overall contribution to the total RNA pool compared to the less numerous myeloid cells, which have substantially more RNA molecules per cell (Suppl. Fig. 2A). Thus, the ‘bulk’ RNA obtained from tumor samples is not dominated by RNA originating from T cells, suggesting that other cell types also contribute to the 3’ UTR shortening that was previously described in tumors. We therefore carried out a cell type-specific analysis of 3’ UTR usage in tumors relative to matched controls.

### A PAS-agnostic approach to quantify 3’ UTR shortening and APA events

A few approaches have been proposed for assessing APA in scRNA-seq data sets (Patrick et al., 2020; Shulman & Elkon, 2019; Wu et al., 2020). However, their robustness with respect to the sparsity of the data and the incompleteness of PAS annotation has not been checked (Ye et al., 2020). Thus, we developed a novel approach (single cell analysis of 3’ untranslated region lengths, SCUREL) (Figure 2A), specifically designed to circumvent these issues and implemented in a Snakemake (Koster & Rahmann, 2012) workflow. SCUREL enables two different comparisons of 3’ UTR length: between two different cell types in a data set (“cell type” mode), or for the same cell type between two different conditions (e.g. tumor and matched control tissue, “condition” mode). We frame the detection of changes in 3’ UTR length between two groups of cells as a problem of identifying the cell group from which the reads originated by inspecting the positions where the reads map in the terminal exons (TEs). That is, read 3’ ends are tabulated and the cumulative coverage along individual TEs is calculated and normalized (Fig. 2B). Then, analyzing each TE individually, we record the fraction of reads from the two cell groups that map within an extending window of the TE starting from the 3’ end (Fig. 2C). This yields a curve in the plane defined by the proportions of reads in the two cell groups, which is similar to a receiver operating characteristic (ROC). The area under this curve (AUC) indicates the similarity of TE length between the compared cell groups. The curve is anchored at coordinates (0,0), corresponding to the end of the TE, where no reads have been observed yet, and (1,1), corresponding to the start of the TE, where all reads from the TE have been accounted for. If the coverage of a TE by read 3’ ends were similar between the two groups of cells and thus the cell group cannot be identified from the position of the reads, the curve would trace the diagonal line. Deviations above the diagonal indicate higher coverage of the distal region of the TE in the cell group represented on the y-axis, while deviations below the diagonal line indicate higher coverage of the distal TE region in the cell group represented on the x-axis. When the number of read mapping to a given TE is small, the curve will show discrete jumps of 1/*n* step size (where *n* is the number of reads mapping to the TE), as individual reads are encountered along the TE. This could lead to AUC values that deviate strongly from the 0.5 value expected under the assumption of similar coverage in the two cell groups. To avoid false positives that are caused by these finite sampling effects, we constructed a background coverage data set by randomizing the labels indicating the cell group from which each read originated. This preserves the depth of coverage of each TE in each group of cells while randomizing the location of each read, thus allowing us to determine changes in 3’ UTR length that cannot be explained by the sparsity of the data. For considerations of efficiency, we carried out the randomization once, and used the information from TEs with similar average coverage to detect significant AUC values. That is, the distribution of AUC values being wider for TEs with low coverage (in counts per million, CPM) compared to TEs with high coverage (Fig. 2D), we binned TEs by the average coverage in the two cell groups (in log(mean CPM)) and within each of the 20 bins, we used the 1% quantile of the randomized read data as the threshold for significant AUC values. Finally, noting that in some cases the difference in TE exon was small and unlikely to be due to APA, we selected only those TEs for which the read 3’ ends span a sufficiently large distance. That is, we calculated the interquartile range (IQR) of read 3’end positions and, if the union of these intervals for the two cell clusters that were analyzed was larger than 200 nucleotides, we considered the range of 3’ end variation sufficient to be indicative of APA (Fig. 2D).

**Figure 2.**
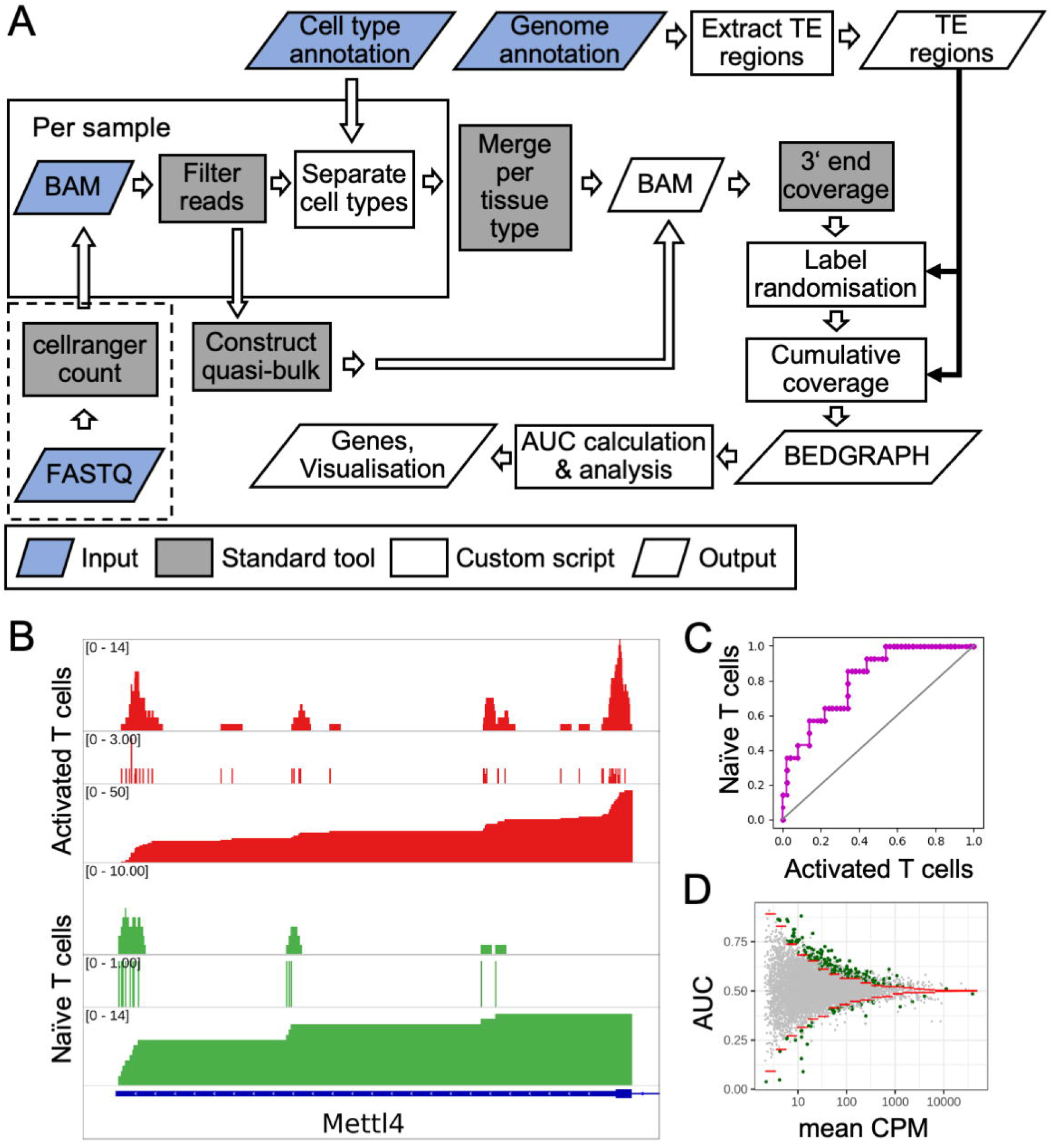
Overview of SCUREL. **A.** Schematic representation of the workflow for detecting significant changes in 3’ UTR length between two cell populations. Input data (blue) consist of mapped reads from *cellranger count* and a table of annotated cell barcodes. The genome annotation is used to extract TEs, their cumulative 3’ end coverage in the two cell groups yielding the AUC value, which we used as a measure of APA. Dashed box: Alternative start of the workflow, from scRNA-seq reads in FASTQ format. The cell type annotation is done semi-automatically, based on marker gene expression (see Methods). **B.** Cumulative 3’ end coverage of the TE of mouse *Mettl4* gene in activated (red) and naive (green) T cells from the (Pace et al., 2018) study. For each cell type, the first track shows the read coverage along the TE, the second track the location of read 3’ ends and the third track the reverse cumulative of the 3’ end coverage. The gene is on the negative strand of the chromosome. **C.** Summary of the cumulative 3’ end read distribution along the TE of *Mettl4* in activated versus naive T cells, from the 3’ (at 0,0) to the 5’ (at 1,1) end. Points correspond to individual nucleotides of the TE where 3’ end reads are observed. The upwards deviation of the curve relative to the diagonal line indicates higher coverage of the distal region of the TE in naive T cells, quantified by the AUC value of 0.582. **D.** Distribution of AUC values as a function of log10(mean CPM) per TE in the mouse T cell activation data set (Pace et al., 2018). 9’099 TEs are represented, 218 showing significant shortening and 43 TEs significant lengthening (green points) attributed to APA.

### SCUREL detects 3’ UTR length changes in previously characterized systems

To validate our approach, we analyzed the dynamics of 3’ UTR length in two well-characterized cellular systems, namely T cell activation, where 3’ UTRs become shorter, and sperm cell development, where the 3’ UTRs are known to become longer. Furthermore, we compared our results with those generated on these data sets by the previously published scAPA method (Shulman & Elkon, 2019).

We annotated the mouse T cell scRNA-seq data (Pace et al., 2018) with *Seurat*, obtaining 1605 activated and 1535 naïve T cells (Figure 3A), with 5.8 and 1.8 million reads mapped to TEs, respectively. Applying SCUREL, we identified 261 TEs whose length changed significantly upon T cell activation, of which 218 (84%) became shorter (Figure 3B). These results recapitulate those obtained from bulk RNA sequencing in a similar system (A. R. Gruber et al., 2014). Applying the previously published scAPA method (Shulman & Elkon, 2019) (see Methods) we only obtained 14 TEs with a significant length change, 12 of which (85%) became shorter (Figure 3C). ⅔ of the scAPA-identified targets (8 of 12 TEs) were also identified by our method, while the 4 cases missed by SCUREL involved either very small TE length changes (3 cases) or a difference in the annotation of the TE, because scAPA also quantifies PAS downstream of annotated TEs. In contrast, inspection of 9 randomly chosen TEs identified only by SCUREL indicated that they correspond to genes with relatively low expression, which are overlooked by scAPA (Suppl. Fig. 3). Examples of TEs from each of these categories are shown in Fig. 3G.

**Figure 3:**
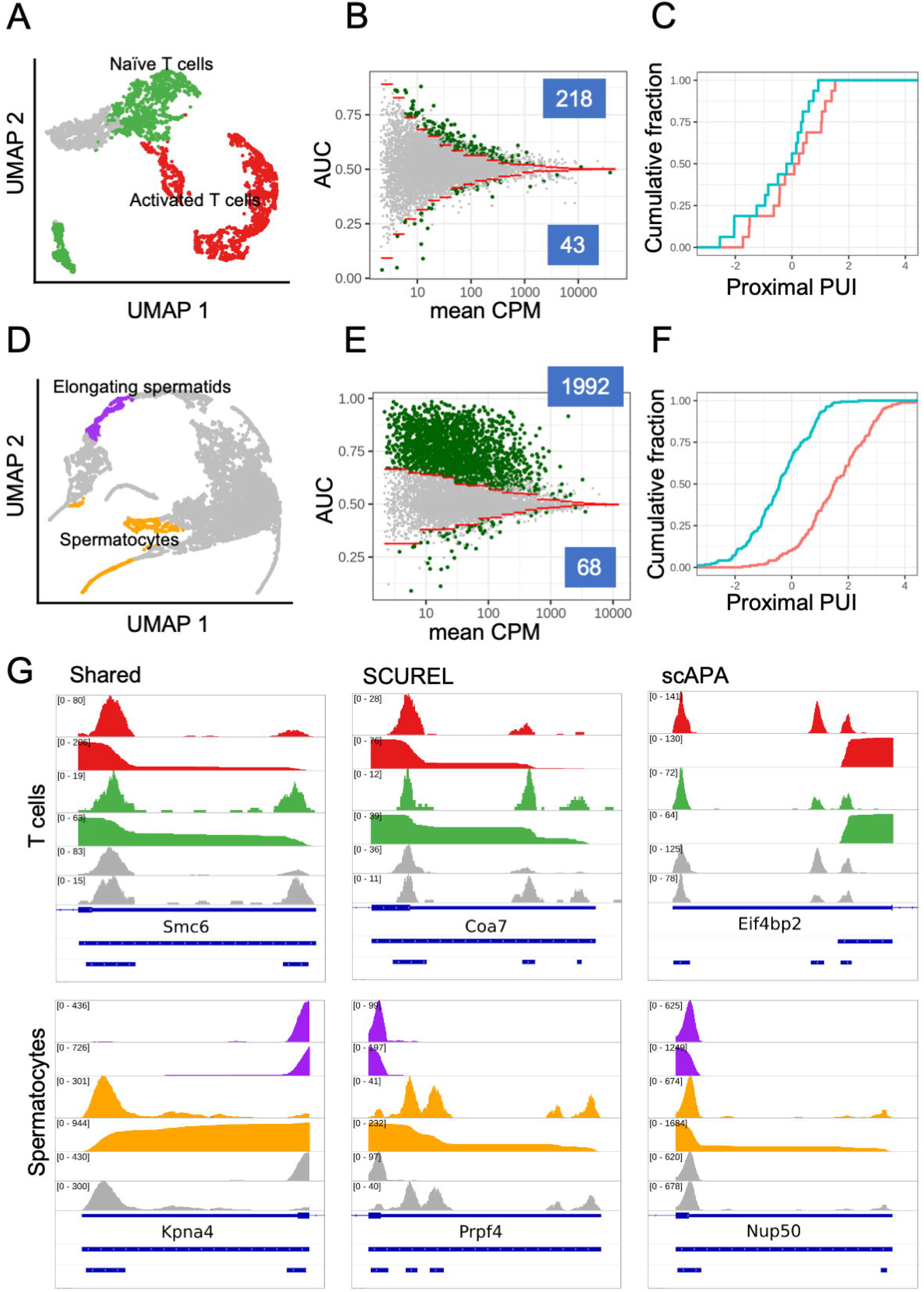
Analysis of APA in T cell activation and spermatogenesis. **A.** UMAP projection of the T cell activation dataset (Pace et al., 2018) showing activated (red), naive (green) and unassigned (grey) T cells. **B.** Scatter plot of AUC in function of log10(mean CPM) for 9’099 TEs. The 1% quantiles (red lines) of the distributions obtained from the randomized dataset were used to identify TEs whose length changed significantly. AUC values > 0.5 indicate shorter 3’ UTRs in activated T cells. TEs whose length changes were attributed to APA based on the span of the read 3’ ends (see Methods) are shown in green. **C.** Cumulative distribution of proximal peak usage index (proximal PUI) for genes deemed by scAPA to undergo significant 3’ UTR length changes. Activated T cells (red) generally have higher proximal PUI compared to naive T cells (blue), indicating 3’UTR shortening in activated T cells. **D**. UMAP projection of the spermatogenesis dataset (Lukassen et al., 2018), with highlighted elongating spermatids (purple) and spermatocytes (orange). **E.** Scatter plot of AUC in function of log10(mean CPM) for 7’875 TEs (see panel B for details). AUC values > 0.5 indicate longer 3’ UTRs in spermatocytes. **F.** As in C, but comparing elongating spermatids (red) with spermatocytes (blue). **G.** Examples of genes deemed to exhibit significant change in 3’ UTR length by both methods (left), by SCUREL only (middle) or by scAPA only (right). For each example, the tracks are: read coverage and cumulative distribution in the two conditions (activated - red - and resting - green - T cells for T cell examples, elongating spermatids - purple - and spermatocytes - orange - for the spermatogenesis examples, followed by coverage tracks from scAPA for the same two conditions in grey. The three blue tracks on the bottom denote in order, the Refseq annotation of the gene, the TE region analyzed in SCUREL and the peaks identified by scAPA.

We carried out a similar analysis on a mouse spermatogenesis dataset (Lukassen et al., 2018), as it is well known that 3’ UTRs become progressively longer during the maturation of germ cells to elongating, condensing, round spermatids and finally spermatocytes. We used the markers described in the original publication (Lukassen et al., 2018) to annotate 386 elongating spermatids (ES) and 667 spermatocytes (SC), with 8 and 12 million reads in the TE regions, respectively (Figure 3D). Applying SCUREL, we found 2’060 TEs whose length changed significantly from ES to SCs, almost all of which (1’992, 97%) became longer (Figure 3E). scAPA yielded a similar proportion of shortened TEs (but fewer in absolute number), 96% (165 of 171 significant APA events, Figure 3F). As in the case of T cells, most of the scAPA-identified TEs were also found by our method (146 of 165 TEs), while TE annotation and small changes in PAS usage accounted for the cases that were unique to scAPA. Inspection of 9 randomly chosen TEs identified only by SCUREL indicated that they correspond to genes with relatively low expression or exclusively express one PAS or the other (Suppl. Fig. 4).

### Genes involved in protein metabolism are targets of 3’ UTR shortening in lung cancer cells

Having established that our method reproduces previously reported patterns of 3’ UTR length change in physiological settings, we then turned to the question of whether 3’ UTRs are also different in lung cancer cells compared to their non-malignant counterpart, the alveolar epithelial cells. We identified 1’330 TEs that were shorter in the 3’607 cancer compared to the 851 alveolar cells in the Lambrechts dataset (with 22 and 3.7 million reads in TEs respectively), representing 98% of 1’357 significant events (Figure 4A, top). Similarly, we identified 188 shortened TEs from the Laughney dataset of 489 cancer and 292 alveolar cells (with 6 and 1.3 million reads in TEs respectively), representing 85% of 219 significant events (Figure 4A, bottom). While much fewer events were found in the Laughney data set, the majority (105 of 188 TEs, 56%) were shared with the Lambrechts dataset. To determine whether specific biological processes are subject to APA-dependent regulation in cancer cells, we submitted the set of 105 shared genes to functional analysis via the STRING web server (Szklarczyk et al., 2019). This revealed that the corresponding proteins are associated with membranes, vesicles and granules (Figure 4B,C). Interestingly, these APA targets cover the entire lifecycle of membrane and secreted proteins, from synthesis (i.e. translation initiation factors and ribosomal proteins), to traffic into the ER (e.g. *SSR1, SPCS3, SEC63*) and Golgi (e.g. *TRAPPC3, KDELR2*), to proteasome-mediated degradation (*PSMD12*). Some of the APA targets are surface receptors with well-known involvement in cancers (*CD44, CD47* and *CD59*). These results indicate that APA contributes to the orchestration of protein metabolism and traffic in cancer cells. Examples of TEs from Figure 4B are shown in Figure 4D.

**Figure 4:**
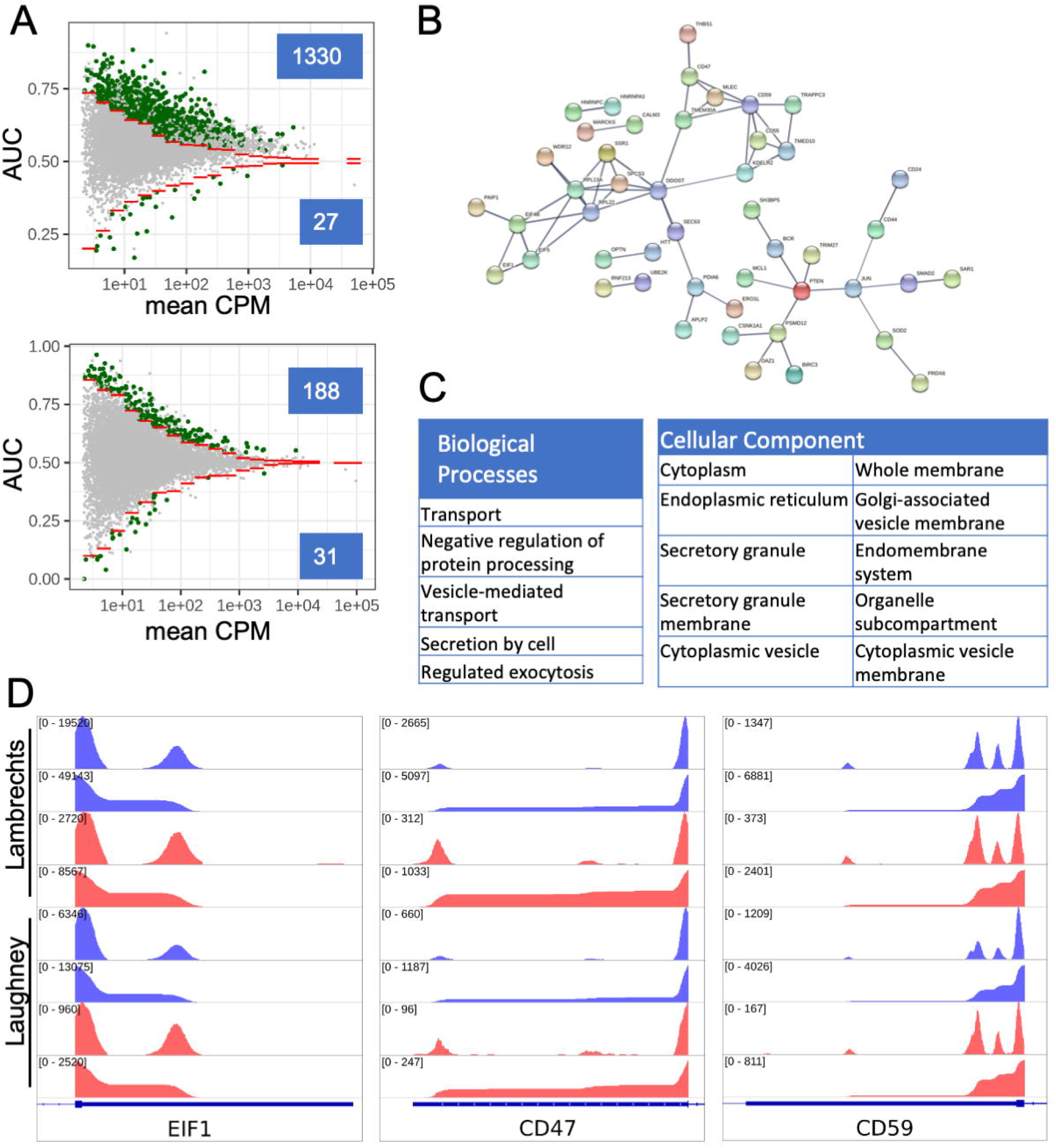
APA in lung adenocarcinoma cells. **A.** Scatter plot of AUC in function of log10(mean CPM) for cancer and alveolar cells in the Lambrechts (top) and Laughney (bottom) datasets. TEs with significant APA-induced length changes are highlighted in green (numbers shown in insets). **B.** The interaction network (from the STRING web server) of proteins whose transcripts undergo 3’UTR shortening in both datasets. **C.** Functional enrichment analysis for genes whose TEs undergo shortening in cancer cells. Shown are the top 10 GO biological process terms (sorted by the false discovery rate, FDR). Analysis was performed with STRING web server, using as background the set of genes found to be expressed in the lung samples. **D** Read coverage along TEs for a few example genes from panel B (*EIF1, CD44* and *CD59*).Each panel shows four tracks per data set, blue: cancer cells, red: alveolar cells, coverage of the TE by reads (top track) and the cumulative coverage of the TE by read 3’ ends (bottom track). In all cases, the 3’ UTRs are shorter in cancer compared to alveolar cells.

### Conserved targets of 3’ UTR shortening in individual cell types

The next question we wanted to answer is whether 3’ UTR shortening affects all cells in the tumor environment, or it is rather restricted to specific cell types. We thus carried out the SCUREL analysis for each individual cell type for which we had at least ~20 cells in each data set, comparing TE lengths between cells of the same type, from the tumor sample and matched control sample. We found many more TEs becoming significantly shorter than longer (Fig. 5A-B), across almost all cell types and in both data sets. This is summarized in Fig. 5C, which shows that the proportion of shortened among significantly changed TEs is almost always greater than 0.5. By grouping all reads from the tumors and from matched control samples, respectively, we also recapitulated the result of previous ‘bulk’ RNA-seq data analyses (Fig. 5D). Thus, 3’ UTR shortening is not restricted to a specific cell type, but seems to generally take place in all cell types, associated with the tumor environment.

**Figure 5.**
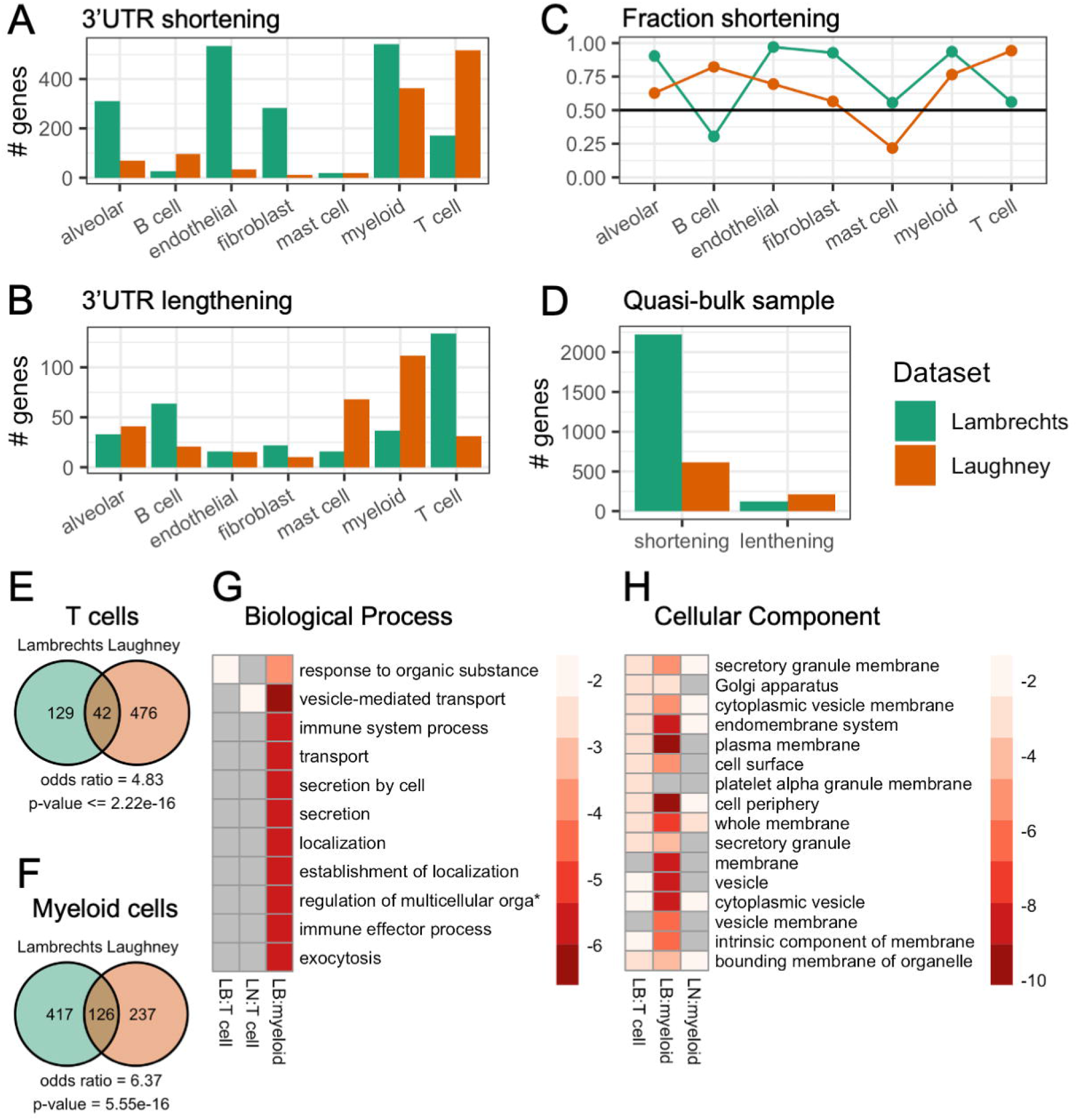
APA events in individual cell types. **A.** Number of genes with APA-associated 3’UTR shortening in the Lambrechts (green) and Laughney (orange) data sets. **B.** Number of genes with APA-associated 3’UTR lengthening, same colors as A. **C.** Fraction of 3’UTR shortening events in individual cell types, among all significant events. **D.** Number of genes whose TEs undergo significant length change in quasi-bulk samples, shortening and lengthening events being shown separately. **E.** Venn diagram of TE shortening events in T cells from the two studies. Calculation of odds ratio and p-value of overlap with hypergeometric distribution (see Methods). **F.** Similar for myeloid cells. **G.** Biological process enrichment for TEs undergoing significant shortening in T cells and myeloid cells from the Lambrechts (LB) and Laughney (LN) studies. No process was specifically enriched in myeloid cells from the Laughney dataset. Plot generated with *pheatmap* (v 1.0.12). **H** Cellular component enrichment for TEs undergoing significant shortening in T cells and myeloid cells from the two studies. No component was specifically enriched in T cells from the Laughney dataset. Plot generated as in G.

Moreover, in spite of the differences between the studies, there was a highly significant overlap between the targets of TE shortening in individual cell types (Fig. 5E-F). To gain further insight into the processes that may be regulated by APA, we submitted the intersection sets of genes exhibiting TE shortening in T lymphocytes and myeloid cells in these studies to functional enrichment analysis. We found significant enrichments especially in cellular components such as membranes, vesicles and granules (Fig. 5G-H), similar to what we observed in cancer cells.

### Variability in 3’ UTR shortening among individuals

Finally, we asked to what extent are the targets of 3’ UTR shortening similar across patients. To answer this question, we analyzed individually the cells obtained from three patients in the Lambrechts study. Interestingly, in spite of the similar histopathological classification of the samples, one of the three samples was markedly different from the others, not exhibiting any tendency towards 3’ UTR shortening (Fig. 6A-D). The other two samples showed highly significant overlaps between shortened 3’ UTRs in different cell types (Fig. 6E). Analysis of biological process enrichment in individual cell types based on the genes targeted in both of these patients reinforced the concept that transport processes are affected in multiple cell types (Fig. 6F). It also provided further granularity. For example, leukocyte activation and secretion are terms enriched in the myeloid cell data, whereas metabolic processes are enriched in T cells, interaction with immune cells in endothelial cells and interaction with endothelial cells and angiogenesis in fibroblasts. Altogether these data demonstrate the power of SCUREL identifying changes in APA-related changes in 3’ UTR length, revealing common functional themes, in spite of substantial variability between samples. A complete table of genes with significant 3’UTR shortening across all LUAD comparisons we conducted is available in Suppl. Table 1. The data further indicate that protein transport processes and intercellular communication are preferential targets of APA across multiple cell types.

**Figure 6.**
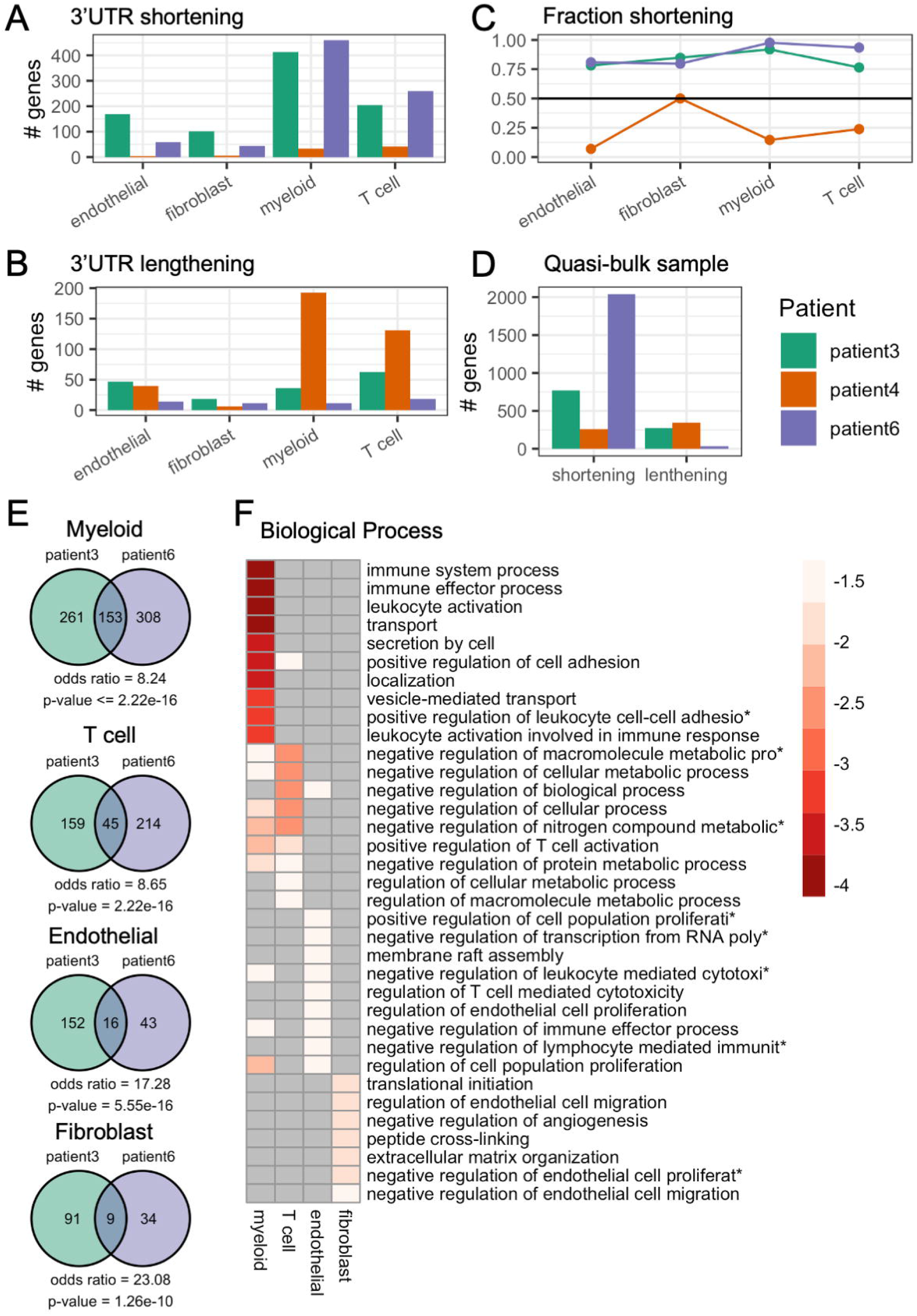
APA events in individual cell types from individual patients. **A.** Number of genes with 3’UTR shortening inferred from patient 3 (green), patient 4 (orange) and patient 6 (purple) samples from the Lambrechts dataset. **B.** Number of genes with 3’UTR lengthening, same colors as A. **C.** Fraction of 3’UTR shortening events in individual cell types, among all significant events. **D.** Number of genes whose TEs undergo significant length change in quasi-bulk samples, shortening and lengthening events being shown separately. **E.** Venn diagrams of significantly shortened TEs in myeloid, T, endothelial and fibroblast cells from tumor relative to matched control samples from distinct patients. Calculation of odds ratio and p-value of overlap with hypergeometric distribution (see Methods). **F.** Biological process enrichment for TEs found to be shorter in cancer compared to matched control cells of individual cell types, from patient 3 and patient 6. Plot generated with *pheatmap* (v 1.0.12).

## Discussion

The remodeling of gene expression in cancers involves, among other processes, alternative polyadenylation. A tendency toward 3’ UTR shortening has been generally observed, though to different extents, in virtually all studied cancers (Schmidt et al., 2018; Xia et al., 2014). Whether this is the result of changes in the cell type composition of the tissue or to cancer-related changes in functionality in all cell types has not been investigated so far. We set out to answer this question, taking advantage of single cell sequencing data sets obtained from human lung adenocarcinoma. As the sparsity of the scRNA-seq data poses some challenges (Lähnemann et al., 2020) we sought two distinct studies that used the same sequencing platform, to identify shared patterns of variation. Furthermore, we developed an approach that controls for both imperfect annotation of transcript isoforms and low read coverage in scRNA-seq.

Comparing data from cells of the same type, but originating either from tumor samples or from matched control tissue, we found similar tendencies towards 3’ UTR shortening in the tumor environment for most cell types. Furthermore, the proteins encoded by the transcripts that are affected in various cell types cluster into specific functional classes, specifically the synthesis, traffic, secretion and degradation of proteins. This implicates APA in the regulation of protein metabolism and the organization of subcellular structure.

Initial studies that described the phenomenon of 3’ UTR shortening in T cells and cancer cells proposed a role in the regulation of protein levels, as short 3’ UTR isoforms are more stable than those with long 3’ UTRs (Mayr & Bartel, 2009; Sandberg et al., 2008). However, when the decay rates of 3’ UTR isoforms were measured, they turned out to be rather similar (A. R. Gruber et al., 2014; Spies et al., 2013), leaving open the question of functional differences between 3’ UTR isoforms (Mayr, 2018). More recent work uncovered additional layers of 3’ UTR-mediated regulation. For example, a role of 3’ UTRs in the localization of the translated protein (UDPL) has been described for a number of membrane proteins, including the immunoglobulin family member CD47, whose localization to the cell membrane protects host cells from phagocytosis by macrophages (Berkovits & Mayr, 2015). Interestingly, *CD47* is a conserved APA target in both LUAD datasets that we analyzed here, its 3’ UTR becoming shorter in cancer cells compared to lung alveolar cancer cells. This would predict decreased localization of CD47 to the surface of cancer cells, making them more susceptible to apoptosis compared to normal alveolar cells. This may explain why increased levels of CD47 are associated with increased cancer-free survival of patients with lung cancers (kmplot.com, (Nagy et al., 2021)). It will be very interesting to apply methods for simultaneous profiling of protein and mRNA expression in single cells (Stoeckius et al., 2017) to better understand the interplay between APA, gene expression, and membrane localization of CD47 in cancers.

The concept that 3’ UTR shortening is associated with proliferative states was challenged in a recent study that instead demonstrated its association with the secretion of proteins, both in trophoblast and in plasma cells (Cheng et al., 2020). Our data fully support this notion, extending the data to cancer cells as well as T lymphocytes and myeloid cells. As the protein production apparatus is present in all cells, APA is a well-suited mechanism for fine-tuning the expression of various components in a cell type and cell state-dependent manner (Lianoglou et al., 2013). Associating APA with protein metabolism rather than cell proliferation makes the question of its upstream regulation ever more puzzling because the shortening of 3’ UTRs in proliferating cells has been attributed to an increased expression of 3’ end processing factors mediated by cell cycle-associated E2F transcription factors (Elkon et al., 2012). It will be interesting to revisit this issue in a system where the increased protein production and secretion can be decoupled from cell proliferation, as the B cell maturation system (Cheng et al., 2020).

In conclusion, among the many applications of scRNA-seq, analysis of cell type-dependent polyadenylation reveals the relevance of APA as a general mechanism for regulating the metabolism and traffic of proteins within cells. With SCUREL we provide a robust method for detecting changes in 3’ UTR length for even low-expression genes between cell types, in a manner that does not rely on accurate PAS annotation.

## Materials and Methods

### Datasets

#### Lung cancer samples

Lung adenocarcinoma (LUAD) and matched control samples were downloaded from the GEO database (Barrett et al., 2013), based on the accession numbers in the original publications. Specifically, from the (Lambrechts et al., 2018; Szklarczyk et al., 2019) data set we used the LUAD samples listed in Table 1 of the original publication (corresponding to patients 3, 4 and 6, 3 tumor samples and one matched control sample for each patient). scRNA-seq data (ArrayExpress (Athar et al., 2019) accession numbers E-MTAB-6149 and E-MTAB-6653) were generated in this study with the 10X Genomics Single Cell 3’ V2 protocol. From the (Laughney et al., 2020) study we also used LUAD and matched control samples, which originated from 3 donors. These samples were also generated with the 10X Genomics Single Cell 3’ V2 protocol (accession number GSE123904).

#### Mouse testis samples

scRNA-seq data from the testes of two 8-week old C57BL/6J mice (Lukassen et al., 2018) were downloaded from the GEO database (accession number GSE104556).

#### Mouse T cell samples

scRNA-seq data of FACS sorted T cells from the lymph nodes and spleen of C57BL/6J mice, three infected with OVA-expressing *Lysteria monocytogenes* and one naive (Pace et al., 2018) were downloaded from the GEO database (accession number GSE106268).

### Execution of scAPA

scAPA (Shulman & Elkon, 2019) was downloaded from the github repository and executed with the same genome sequence that was used throughout the study. For compatibility, the “chr” prefix in the chromosome names was removed. The lengths of the chromosomes were obtained with *samtools faidx*. The *homer* software (v4.11.1) required by the scAPA package was manually downloaded from http://homer.ucsd.edu/homer/. We collected all other requirements specified on scAPA github page in a conda environment. The removal of duplicate reads was done by adjusting the existing *umi_tools dedup* command in *scAPA.shell.script.R* for 10X Genomics, using the following options “ --per-gene”, “ -according to the protocolwgene-tag=GX”, “ --per-cell “. This was necessary because according to the protocol, one RNA fragment could result in reads that do not map at identical positions.

### Extraction of terminal exons

Terminal exons were obtained from the RefSeq genome annotations (gff), GRCm38.p6 for mouse and GRCh38.p13 for human, with a custom script, as follows. Chromosome names from the RefSeq assembly were converted to ENSEMBL-type names based on the accompanying ‘assembly_report.txt’ file. Only autosomes, allosomes and mitochondrial DNA were retained. Based on the genome annotation file, protein-coding and long non-coding transcripts were retained, while model transcripts (‘Gnomon’ prediction; accession prefixes XM_, XR_, and XP_) were discarded. From this transcript set, the 3’-most exons (i.e. terminal exons, TEs) were retrieved. Overlapping TEs on the same chromosome strand were clustered with *intervaltree* (v3.0.2; python package) and from each cluster, the longest exon was kept. The resulting set of TEs was sorted by chromosome and start position and saved to a BED-formatted file. TE IDs were converted to gene names with *biomaRt* (v 2.46.3) using the ensembl BioMart database. Duplicate gene names were discarded.

### Processing of scRNA-seq reads

The workflow can start from mapped reads in *cellranger*-compatible format, a file with cell barcode-to-cell type annotation and a genome annotation file. Alternatively, the *cellranger count* function can be used to map reads from FASTQ input data. Reads from the FASTQ files were mapped with the function *count* from the *cellranger* (v5.0.0) package to the reference human genome GRCh38-3.0.0 sequence obtained directly from *10X genomics* website. This genome is a modified version of the GRCh38 genome, compatible with the cellranger analysis pipeline. Reads are also aligned to the transcriptome. In this step, cell barcodes and UMIs correction also takes place. Aligned reads (BAM) with mapping quality (MAPQ) scores > 30 were selected with *samtools* (v1.12, (Li et al., 2009)). Reads without a cell barcode “CB” tag were removed with *samtools view*, as were duplicated reads using *umi_tools dedup* (v1.1.1, (Smith et al., 2017)). The mapped reads are filtered, deduplicated and grouped by cell type in the “cell type” mode or by cell type and tissue of origin in the “condition” mode. In the latter case, quasi-bulk samples are also constructed from the filtered reads that come from individual conditions.

### Cell type annotation

The annotation of cell types in all datasets was carried out with the approach described in (Lambrechts et al., 2018). Filtered data (so as to remove artifacts such as empty droplets) consisting of cellular barcodes and count matrices from individual data sets were loaded in R (v4.0.3) with *Read10X* (from *Seurat* v3.2.3 (Butler et al., 2018)), and Seurat objects were created with *CreateSeuratObject*. For the lung cancer datasets, cells with < 201 Unique Molecular Identifiers (UMIs), with < 101 or > 6000 genes or with > 10% UMIs from mitochondrial genes (which may indicate apoptotic or damaged cells) were removed. For all datasets, genes with zero variance across all cells (i.e. sum = 0) were discarded. The gene expression counts for each cell were log-normalised with *NormalizeData* with a default scale factor of 10’000. In Seurat, 2’000-2’500 most variable genes are used to cluster the cells. Here we used the 2’192 variable most variable genes, as in (Lambrechts et al., 2018). These were selected with *FindVariableFeatures*, with normalised expression between 0.125 and 3, and a quantile-normalised variance exceeding 0.5 for lung cancer and mouse T cell samples, and normalised expression between 0.1 and 8 for mouse testis samples. Gene expression levels were then centered and scaled across all cells. After Principal Component Analysis (PCA) on the most variable genes, the number of relevant dimensions *n* for each data set was determined by assessing the variance explained by individual Principal Components (PC) with *ElbowPlot* from Seurat. UMAP (McInnes et al., 2018) was used to visualize the data projected on the *n* dimensions. For T cell activation and LUAD samples, batch correction and data integration were performed with *harmony* (v1.0) (Korsunsky et al., 2019). *Harmony* was run on the first 30 PCs and set to group by dataset. The transformed data set was used for downstream analysis (i.e. clustering of cells, visualization in 2D).

Various Seurat functions were used to identify the cell type of individual cells. Cells were clustered using the Shared Nearest Neighbor (SNN) algorithm, which aims to optimize modularity. First, *FindNeighbors* was executed using the first *n* dimensions from PCA or *harmony* and with otherwise default settings (k = 20). Then, *FindClusters* with resolution parameter 0.6 for LUAD, 0.2 for T cells and 0.3 for spermatocytes was run, so as to retrieve a number of clusters similar to those in the original publications. The expression of cell type markers in each cluster was assessed with *FindAllMarkers*. This function finds genes that are differentially expressed between cells from one cluster and all other cells, by applying a Wilcoxon Rank Sum test on the log-normalized expression. Individual clusters were downsampled to the number of cells in the smallest cluster or to at least 100 cells. Only genes expressed in a minimum of 10% of the cells in either population and with a log (base *e*) -fold-change of at least 0.25 (default values in *Seurat*) were tested. Markers with adjusted p-value < 0.01 were considered significant and those with higher expression in the selected cluster were considered as potential markers for that cell cluster. For each cluster we counted the number of significant markers that matched known cell type markers (Table 1) and assigned the cell type to be the one for which a proportion of > 0.6 of known markers were specifically expressed in the cell cluster. Generally, this assignment was unambiguous, and when it was not, the cell type assignment was done manually, taking into account the adjusted p-value and average log-fold-change of all considered marker genes as well as the cell type annotation from the Suppl. Table 3 of (Lambrechts et al., 2018), which contains additional cell type markers. At least 3 marker genes were required to assign a cluster to the corresponding cell type, except for cancer cells that were annotated only based on the expression of EPCAM.

### Assessing 3’ UTR length differences with the AUC measure

To assess changes in 3’ UTR length between groups of cells we used the following approach. For simplicity, the analysis is carried out for terminal exons (TEs) rather than 3’ UTRs, as 3’ UTRs are generally contained in TEs, covering almost the entire length of the TEs. We started from the BAM files of mapped reads from two groups of cells. We computed the 3’ end coverage of individual TEs per strand with *bedtools genomecov* and parameter “-bga”. The BED file with read 3’ end positions was used to obtain the normalized reverse cumulative coverage of individual TEs, i.e. starting at the TE 3’ end and ending at the 5’ most nucleotide. Individual TEs were traversed from the end to the beginning, recording the reverse cumulative coverage in the two groups of cells as a function of position. The area under the resulting curve (AUC) was then calculated. An AUC of 0.5 corresponds to identical position-dependent coverage of the TE by 3’ end reads in the two groups of cells, i.e. no difference in TE length. An AUC value of 1 corresponds to all the 3’ end reads from the group of cells indicated on the y-axis being clustered at the end of the TE, before any reads from the other group are observed, thus the TEs are longest in this group of cells. Vice versa, an AUC value of 0 corresponds to all the 3’ end reads from the group indicated on x-axis are observed before any reads of the other group, thus the 3’UTRs are longest in this group of cells.

If the read coverage of a TE is very sparse, the curve representing the coverage in the two cell groups will not be smooth, but rather change in steps of 1/*n* where *n* is the number of reads mapping to the TE; deviations from the diagonal line of identical coverage in the two groups will be common, due to the stochastic sampling of the reads. To mitigate this effect and identify TEs whose coverage cannot be explained by stochastic sampling of low-expression genes we generated a background dataset, in which we randomized the cell group label of the reads. This procedure preserves the number of reads obtained in each TE in each group, but randomizes their position in the TE.

Finally, we identified TEs with AUCs indicating significant shifts in PAS usage. For this, we extracted TEs with a normalized read count (CPM) >= 2 in both cell groups, roughly corresponding to TEs with at least one count in each of the groups. As AUC values depend on the overall expression of the TE, we used an expression-dependent AUC cutoff to identify the TEs significantly changing length. This corresponded to the two-tailed 1% quantile of the background distribution in each of the 20 equal-sized log(mean expression between cell groups) bins, smoothened using the median over a running window of 5 values. Finally, to ensure that the change in read coverage was due to APA, we only retained significantly changed TEs for which the union of the interquartile range of TE positions that were covered by 3’ end reads in the two samples spanned at least 200 nucleotides.

### Analysis of overlaps between data sets

We used a sample-specific background for the calculation of the probability of overlap of genes and for the pathway enrichment analysis carried out on the STRING web server. All TEs considered in the AUC analysis, i.e. TEs with CPM >= 2, in each sample were combined and the unique set of TEs was used as background. In particular, for the cell type analysis of the Lambrechts dataset, we used the cell type-specific union of TEs from patients 3, 4 and 6 and obtained 10’966 genes for myeloid cells, 10’473 for T cells, 11’269 for endothelial cells and 11’857 for fibroblasts. For the cell type analysis of lung cancer datasets, the union of TEs consisted of 10’177 genes in T cells and 9’970 genes in myeloid cells. We used the hypergeometric distribution to calculate the odds ratio and associated p-value of the overlap between gene sets.

### Pathway analysis

The gene symbols for TEs with significant APA events were analyzed via the STRING web server, which provides enriched Gene ontology (GO) terms, KEGG and reactome pathways. As a background gene set for the enrichment analysis we provided the dataset-specific list of expressed genes (CPM >= 2).

### Workflow execution

SCUREL was packaged in Snakemake and can be obtained from https://github.com/zavolanlab/SCUREL.

## Acknowledgements

We thank the tissue donors and the authors of the single cell studies (Lambrechts et al., 2018; Laughney et al., 2020) for providing the data. We thank Erik van Nimwegen for the extensive and fruitful discussions on measures of 3’ UTR shortening. We also thank members of the Zavolan lab and Dr. Sebastian Leidel for constructive comments. We especially acknowledge the help of Arka Banerjee in the cell marker identification and Dr. Jeremie Breda for helpful discussions regarding the AUC measure. We are grateful to the sciCORE team for the maintenance of the computing infrastructure on which the computations were carried out. This work was supported by the Swiss National Science Foundation grant #310030_189063 to M.Z. D.B. is a recipient of a Fellowship for Excellence from the Biozentrum, University of Basel.

## Notes

### Competing Interest Statement

The authors have declared no competing interest.

## References

Athar, A., Füllgrabe, A., George, N., Iqbal, H., Huerta, L., Ali, A., Snow, C., Fonseca, N. A., Petryszak, R., Papatheodorou, I., Sarkans, U., & Brazma, A. (2019). ArrayExpress update - from bulk to single-cell expression data. Nucleic Acids Research, 47(D1), D711–D715.

Barrett, T., Wilhite, S. E., Ledoux, P., Evangelista, C., Kim, I. F., Tomashevsky, M., Marshall, K. A., Phillippy, K. H., Sherman, P. M., Holko, M., Yefanov, A., Lee, H., Zhang, N., Robertson, C. L., Serova, N., Davis, S., & Soboleva, A. (2013). NCBI GEO: archive for functional genomics data sets--update. Nucleic Acids Research, 41(Database issue), D991–D995.

Berkovits, B. D., & Mayr, C. (2015). Alternative 3’ UTRs act as scaffolds to regulate membrane protein localization. Nature, 522(7556), 363–367.

Breda, J., Zavolan, M., & van Nimwegen, E. (2021). Bayesian inference of gene expression states from single-cell RNA-seq data. Nature Biotechnology. https://doi.org/10.1038/s41587-021-00875-x

Butler, A., Hoffman, P., Smibert, P., Papalexi, E., & Satija, R. (2018). Integrating single-cell transcriptomic data across different conditions, technologies, and species. Nature Biotechnology, 36(5), 411–420.

Cheng, L. C., Zheng, D., Baljinnyam, E., Sun, F., Ogami, K., Yeung, P. L., Hoque, M., Lu, C.-W., Manley, J. L., & Tian, B. (2020). Widespread transcript shortening through alternative polyadenylation in secretory cell differentiation. Nature Communications, 11(1), 3182.

Dwyer, D. F., Barrett, N. A., Austen, K. F., & Immunological Genome Project Consortium. (2016). Expression profiling of constitutive mast cells reveals a unique identity within the immune system. Nature Immunology, 17(7), 878–887.

Elkon, R., Drost, J., van Haaften, G., Jenal, M., Schrier, M., Vrielink, J. A. O., & Agami, R. (2012). E2F mediates enhanced alternative polyadenylation in proliferation. Genome Biology, 13(7), R59.

Gruber, A. J., Gypas, F., Riba, A., Schmidt, R., & Zavolan, M. (2018). Terminal exon characterization with TECtool reveals an abundance of cell-specific isoforms. Nature Methods, 15(10), 832–836.

Gruber, A. J., Schmidt, R., Ghosh, S., Martin, G., Gruber, A. R., van Nimwegen, E., & Zavolan, M. (2018). Discovery of physiological and cancer-related regulators of 3’ UTR processing with KAPAC. Genome Biology, 19(1), 44.

Gruber, A. R., Martin, G., Müller, P., Schmidt, A., Gruber, A. J., Gumienny, R., Mittal, N., Jayachandran, R., Pieters, J., Keller, W., van Nimwegen, E., & Zavolan, M. (2014). Global 3’ UTR shortening has a limited effect on protein abundance in proliferating T cells. Nature Communications, 5, 5465.

Ji, Z., & Tian, B. (2009). Reprogramming of 3’ untranslated regions of mRNAs by alternative polyadenylation in generation of pluripotent stem cells from different cell types. PloS One, 4(12), e8419.

Korsunsky, I., Millard, N., Fan, J., Slowikowski, K., Zhang, F., Wei, K., Baglaenko, Y., Brenner, M., Loh, P.-R., & Raychaudhuri, S. (2019). Fast, sensitive and accurate integration of single-cell data with Harmony. Nature Methods, 16(12), 1289–1296.

Koster, J., & Rahmann, S. (2012). Snakemake--a scalable bioinformatics workflow engine. In Bioinformatics (Vol. 28, Issue 19, pp. 2520–2522). https://doi.org/10.1093/bioinformatics/bts480

Lähnemann, D., Köster, J., Szczurek, E., McCarthy, D. J., Hicks, S. C., Robinson, M. D., Vallejos, C. A., Campbell, K. R., Beerenwinkel, N., Mahfouz, A., Pinello, L., Skums, P., Stamatakis, A., Attolini, C. S.-O., Aparicio, S., Baaijens, J., Balvert, M., Barbanson, B. de, Cappuccio, A.,… Schönhuth, A. (2020). Eleven grand challenges in single-cell data science. Genome Biology, 21(1), 31.

Lambrechts, D., Wauters, E., Boeckx, B., Aibar, S., Nittner, D., Burton, O., Bassez, A., Decaluwé, H., Pircher, A., Van den Eynde, K., Weynand, B., Verbeken, E., De Leyn, P., Liston, A., Vansteenkiste, J., Carmeliet, P., Aerts, S., & Thienpont, B. (2018). Phenotype molding of stromal cells in the lung tumor microenvironment. Nature Medicine, 24(8), 1277–1289.

Laughney, A. M., Hu, J., Campbell, N. R., Bakhoum, S. F., Setty, M., Lavallée, V.-P., Xie, Y., Masilionis, I., Carr, A. J., Kottapalli, S., Allaj, V., Mattar, M., Rekhtman, N., Xavier, J. B., Mazutis, L., Poirier, J. T., Rudin, C. M., Pe’er, D., & Massagué, J. (2020). Regenerative lineages and immune-mediated pruning in lung cancer metastasis. Nature Medicine, 26(2), 259–269.

Lee, S., Wei, L., Zhang, B., Goering, R., Majumdar, S., Wen, J., Taliaferro, J. M., & Lai, E. C. (2021). ELAV/Hu RNA binding proteins determine multiple programs of neural alternative splicing. PLoS Genetics, 17(4), e1009439.

Lianoglou, S., Garg, V., Yang, J. L., Leslie, C. S., & Mayr, C. (2013). Ubiquitously transcribed genes use alternative polyadenylation to achieve tissue-specific expression. Genes & Development, 27(21), 2380–2396.

Li, H., Handsaker, B., Wysoker, A., Fennell, T., Ruan, J., Homer, N., Marth, G., Abecasis, G., Durbin, R., & 1000 Genome Project Data Processing Subgroup. (2009). The Sequence Alignment/Map format and SAMtools. Bioinformatics, 25(16), 2078–2079.

Lukassen, S., Bosch, E., Ekici, A. B., & Winterpacht, A. (2018). Characterization of germ cell differentiation in the male mouse through single-cell RNA sequencing. Scientific Reports, 8(1), 6521.

Martin, G., Gruber, A. R., Keller, W., & Zavolan, M. (2012). Genome-wide analysis of pre-mRNA 3’ end processing reveals a decisive role of human cleavage factor I in the regulation of 3’ UTR length. Cell Reports, 1(6), 753–763.

Masuda, A., Kawachi, T., Takeda, J.-I., Ohkawara, B., Ito, M., & Ohno, K. (2020). tRIP-seq reveals repression of premature polyadenylation by co-transcriptional FUS-U1 snRNP assembly. EMBO Reports, 21(5), e49890.

Mayr, C. (2018). What Are 3’ UTRs Doing? Cold Spring Harbor Perspectives in Biology. https://doi.org/10.1101/cshperspect.a034728

Mayr, C., & Bartel, D. P. (2009). Widespread Shortening of 3’UTRs by Alternative Cleavage and Polyadenylation Activates Oncogenes in Cancer Cells. Cell, 138(4), 673–684.

McInnes, L., Healy, J., Saul, N., & Großberger, L. (2018). UMAP: Uniform Manifold Approximation and Projection. In Journal of Open Source Software (Vol. 3, Issue 29, p. 861). https://doi.org/10.21105/joss.00861

Nagy, Á., Munkácsy, G., & Győrffy, B. (2021). Pancancer survival analysis of cancer hallmark genes. Scientific Reports, 11(1), 6047.

Pace, L., Goudot, C., Zueva, E., Gueguen, P., Burgdorf, N., Waterfall, J. J., Quivy, J.-P., Almouzni, G., & Amigorena, S. (2018). The epigenetic control of stemness in CD8 T cell fate commitment. Science, 359(6372), 177–186.

Patrick, R., Humphreys, D. T., Janbandhu, V., Oshlack, A., Ho, J. W. K., Harvey, R. P., & Lo, K. K. (2020). Sierra: discovery of differential transcript usage from polyA-captured single-cell RNA-seq data. Genome Biology, 21(1), 167.

Reyes, A., & Huber, W. (2018). Alternative start and termination sites of transcription drive most transcript isoform differences across human tissues. Nucleic Acids Research, 46(2), 582–592.

Sandberg, R., Neilson, J. R., Sarma, A., Sharp, P. A., & Burge, C. B. (2008). Proliferating Cells Express mRNAs with Shortened 3’ Untranslated Regions and Fewer MicroRNA Target Sites. Science, 320(5883), 1643–1647.

Schmidt, R., Ghosh, S., & Zavolan, M. (2018). The 3’ UTR Landscape in Cancer. In eLS (pp. 1–9). https://doi.org/10.1002/9780470015902.a0027958

Shulman, E. D., & Elkon, R. (2019). Cell-type-specific analysis of alternative polyadenylation using single-cell transcriptomics data. Nucleic Acids Research, 47(19), 10027–10039.

Smith, T., Heger, A., & Sudbery, I. (2017). UMI-tools: modeling sequencing errors in Unique Molecular Identifiers to improve quantification accuracy. Genome Research, 27(3), 491–499.

So, B. R., Di, C., Cai, Z., Venters, C. C., Guo, J., Oh, J.-M., Arai, C., & Dreyfuss, G. (2019). A Complex of U1 snRNP with Cleavage and Polyadenylation Factors Controls Telescripting, Regulating mRNA Transcription in Human Cells. Molecular Cell, 76(4), 590–599.e4.

Spies, N., Burge, C. B., & Bartel, D. P. (2013). 3’ UTR-isoform choice has limited influence on the stability and translational efficiency of most mRNAs in mouse fibroblasts. Genome Research. https://doi.org/10.1101/gr.156919.113

Stoeckius, M., Hafemeister, C., Stephenson, W., Houck-Loomis, B., Chattopadhyay, P. K., Swerdlow, H., Satija, R., & Smibert, P. (2017). Simultaneous epitope and transcriptome measurement in single cells. Nature Methods, 14(9), 865–868.

Szklarczyk, D., Gable, A. L., Lyon, D., Junge, A., Wyder, S., Huerta-Cepas, J., Simonovic, M., Doncheva, N. T., Morris, J. H., Bork, P., Jensen, L. J., & Mering, C. von. (2019). STRING v11: protein-protein association networks with increased coverage, supporting functional discovery in genome-wide experimental datasets. Nucleic Acids Research, 47(D1), D607–D613.

Wu, X., Liu, T., Ye, C., Ye, W., & Ji, G. (2020). scAPAtrap: identification and quantification of alternative polyadenylation sites from single-cell RNA-seq data. Briefings in Bioinformatics. https://doi.org/10.1093/bib/bbaa273

Xia, Z., Donehower, L. A., Cooper, T. A., Neilson, J. R., Wheeler, D. A., Wagner, E. J., & Li, W. (2014). Dynamic analyses of alternative polyadenylation from RNA-seq reveal a 3’-UTR landscape across seven tumour types. Nature Communications, 5, 5274.

Ye, C., Zhou, Q., Wu, X., Yu, C., Ji, G., Saban, D. R., & Li, Q. Q. (2020). scDAPA: detection and visualization of dynamic alternative polyadenylation from single cell RNA-seq data. Bioinformatics, 36(4), 1262–1264.

